# The biggest losers: Habitat isolation deconstructs complex food webs from top to bottom

**DOI:** 10.1101/439190

**Authors:** Remo Ryser, Johanna Häussler, Markus Stark, Ulrich Brose, Björn C. Rall, Christian Guill

## Abstract

Habitat fragmentation is threatening global biodiversity. To date, there is only limited understanding of how the different aspects of habitat fragmentation (habitat loss, number of fragments and isolation) affect species diversity within complex ecological networks such as food webs. Here, we present a dynamic and spatially-explicit food web model which integrates complex food web dynamics at the local scale and species-specific dispersal dynamics at the landscape scale, allowing us to study the interplay of local and spatial processes in metacommunities. We here explore how the number of habitat patches, i.e. the number of fragments, and an increase of habitat isolation, affect the species diversity patterns of complex food webs (*α*-, *β*-, *γ*-diversity). We specifically test whether there is a trophic dependency in the effect of these to factors on species diversity. In our model, habitat isolation is the main driver causing species loss and diversity decline. Our results emphasise that large-bodied consumer species at high trophic positions go extinct faster than smaller species at lower trophic levels, despite being superior dispersers that connect fragmented landscapes better. We attribute the loss of top species to a combined effect of higher biomass loss during dispersal with increasing habitat isolation in general, and the associated energy limitation in highly fragmented landscapes, preventing higher trophic levels to persist. To maintain trophic-complex and species-rich communities calls for effective conservation planning which considers the interdependence of trophic and spatial dynamics as well as the spatial context of a landscape and its energy availability.

## Introduction

Understanding the impact of habitat fragmentation (habitat loss, number of fragments, and isolation) on biodiversity is crucial for ecology and conservation biology [1–3]. A general observation and prediction is that large-bodied predators at high trophic levels which depend on sufficient food supplied by lower trophic levels are most sensitive to fragmentation, and thus, might respond more strongly than species at lower trophic levels [4, 5]. However, most conclusions regarding the effect of fragmentation are based on single species or competitively interacting species (see references within [6–8], but see for example [9–11] for food chains and simple food web motifs). There is thus limited understanding how species embedded in complex food webs with multiple trophic levels respond to habitat fragmentation [4, 12–15], even though these networks are a central organising theme in nature [16, 17].

The stability of complex food webs is, amongst others, determined by the number and strength of trophic interactions [18]. While it is broadly recognised that habitat fragmentation can have substantial impacts on such feeding relationships [19, 20], we lack a comprehensive and mechanistic understanding of how the disruption or loss of these interactions will affect species persistence and food web stability [15, 19, 21, 22]. Assuming that a loss of habitat, a decreasing number of fragments, and increasing isolation of the remaining fragments disrupt or weaken trophic interactions [7], thereby causing species extinctions [15, 20], population and community dynamics might change in unexpected and unpredictable ways. This change in community dynamics might lead to secondary extinctions which potentially cascade through the food web [23, 24].

Habitat loss, i.e. the decrease of total habitable area in the landscape or a reduction in patch size, can limit population sizes and biomass production, which might drive energy-limited species extinct [25, 26] and subsequently entail cascading extinctions [23]. Successful dispersal among habitat patches might prevent local extinctions (spatial rescue effects), and thus, ensure species persistence at the landscape scale [27, 28]. Whether dispersal is successful or not depends, among other factors, on the distance an organism has to travel to reach the next habitat patch and on the quality of the matrix the habitat patches are embedded in (in short: the habitat matrix) [29]. With progressing habitat fragmentation, suitable habitat becomes scarce and the remaining habitat fragments increasingly isolated [3, 30], affecting the dispersal network of a species. As a consequence, organisms have to disperse over longer distances to connect habitat patches, which in turn might increase dispersal mortality and thus promote species extinctions [2]. Also, habitat fragmentation often increases the hostility of the habitat matrix, e.g. due to human land use and landscape degeneration [3, 31, 32]. The increased matrix hostility might further reduce the likelihood of successful dispersal between habitat patches as the movement through a hostile habitat matrix is energy intensive, and thus, population biomass is lost [29, 31]. This loss depends on the distance an organism has to travel and its dispersal ability, i.e. its dispersal range and the energy it can invest into movement. Finally, the detrimental effects of habitat loss and increasing isolation are likely to interact, as dispersal mortality can be expected to have a larger per capita effect when a population is already declining due to decreasing habitat.

In this context, superior dispersers might have an advantage over species with restricted dispersal abilities if the distances between habitat patches expand to a point where dispersal-limited species can no longer connect habitat patches. If this is the case, increasing habitat isolation impedes the ability of organisms to move across a fragmented landscape and prevents spatial rescue effects buffering against local extinctions. Increasing habitat isolation might result in increased extinction rates and ultimately lead to the loss of dispersal-limited species from the regional species pool. As large animal species are, at least up to a certain threshold, faster than smaller ones [33, 34], they should also be able to disperse over longer distances [4, 35, 36]. In fragmented landscapes, this body mass dependent scaling of dispersal range might favour large-bodied consumers such as top predators, and thus, increase top-down pressure resulting in top-down regulated communities.

Empirical evidence and results from previous modelling approaches, however, suggest that species at higher trophic positions are most sensitive to isolation [9, 15, 37–39]. Modelling tri-trophic food chains in a patch-dynamic framework, Liao *et al.* [9, 10] for example, show that increasing habitat fragmentation leads to faster extinctions of species at higher trophic levels, which they ascribe to reduced availability of prey [9]. In the fragmentation experiment by Davies *et al.* [39] on the other hand the observed loss of top species is attributed to the unstable population dynamics of top species under environmental change.

Despite its relevance, a realistic picture and comprehensive understanding of how natural food webs might respond to different aspects of fragmentation such as habitat loss or increasing isolation, and any alteration to the spatial configuration of habitat in general, are lacking. To understand how fragmentation affects the diversity of communities organised in complex food webs requires knowledge of the interplay between their local (trophic) and spatial (dispersal) dynamics. The latter are determined by the number of fragments in the landscape and the distance between them, which can potentially affect the local trophic dynamics. We address this issue using a novel modelling approach which integrates local population dynamics of complex food webs and species-specific dispersal dynamics at the landscape scale (which we hereafter refer to as meta-food-web model, see figure 1 for a conceptual illustration). Our spatially-explicit dynamic meta-food-web model allows us to explore how direct and indirect interactions between species in complex food webs together with spatial processes that connect sub-populations in different habitat patches interact to produce diversity patterns across increasingly fragmented landscapes. Specifically, we ask how the number of fragments and increasing habitat isolation impact the diversity patterns in complex food webs. We further ask which species or trophic groups shape these patterns.

**Figure 1:**
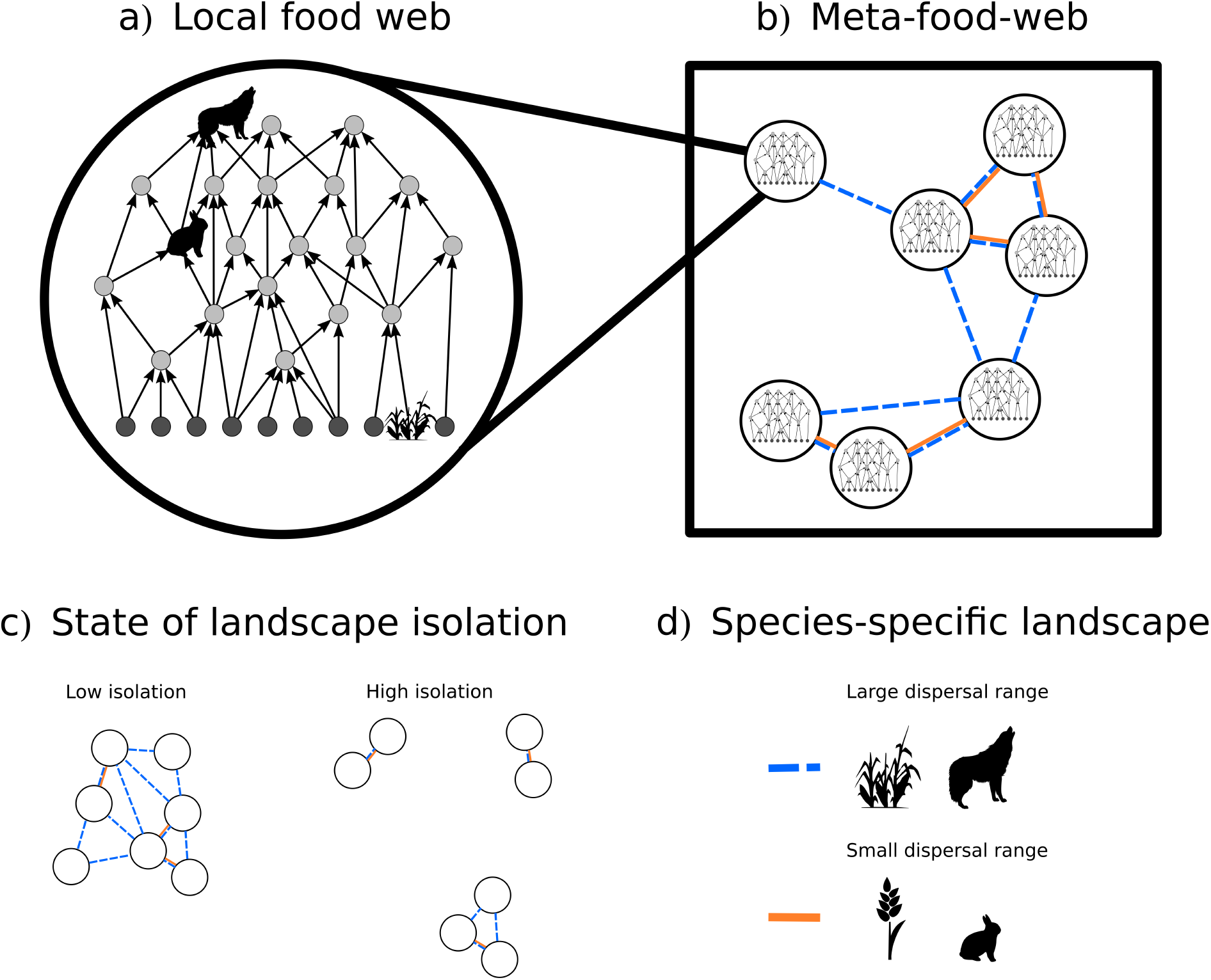
Conceptual illustration of our modelling framework. In our meta-food-web model (b) we link local food web dynamics at the patch level (a) through dynamic and species-specific dispersal at the landscape scale (d). We consider landscapes with identical but randomly distributed habitat patches, i.e. all patches have the same abiotic conditions, and each patch can potentially harbour the full food web. We model fragmented landscapes which differ in the number of habitat patches and the mean distance between patches (c).

Following general observations and predictions, we expect species diversity within complex food webs to decrease along a gradient of isolation. Based on the substantial variation in both dispersal abilities and energy requirements among species and across trophic levels [4, 25, 39], we expect species at different trophic levels to strongly vary in their response to isolation. Specifically, we expect certain trophic groups such as consumer species at lower trophic ranks with limited dispersal abilities or top predators with strong resource constraints to be particularly sensitive to isolation. Additionally, with a larger number of fragments we expect more potential for rescue effects, thus fostering survival. This might especially apply to species with large dispersal ranges, which allow them to connect many habitat patches. We test our expectations using Whittaker’s classical approach of *α*-, *β*-, and *γ*-diversity [40], where *α*- and *γ*-diversity describe species richness at the local (patch) and regional (metacommunity) scale, respectively, and *β*-diversity accounts for compositional differences between local communities.

## Methods

In the following we outline a methods summary, for detailed information on equations and parameters see the methods section in the supplement. We consider a multitrophic metacommunity consisting of 40 species on a varying number of randomly positioned habitat patches (the meta-food-web, figure 1b). All patches have the same abiotic conditions and each patch can potentially harbour the full food web, consisting of 10 basal plant and 30 animal consumer species. The potential feeding links (i.e. who eats whom) are constant over all patches (figure 1a,b) and are as well as the feeding dynamics determined by the allometric food web model by Schneider *et al.* [41]. We use a dynamic bioenergetic model formulated in terms of ordinary differential equations that describe the feeding and dispersal dynamics. The rate of change in biomass density of a species depends on its biomass gain by feeding and immigration and its biomass loss by metabolism, being preyed upon and emigration. We integrate dispersal as species-specific biomass flow between habitat patches (figure 1b,d). Based on empirical observations (e.g. [35]) and previous theoretical frameworks (e.g. [4, 12, 34, 42]), we assume that the maximum dispersal distance of animal species increases with their body mass. As plants are passive dispersers, we model their maximum dispersal distance as random and body mass independent. We model emigration rates as a function of each species’ per capita net growth rate, which is summarising local conditions such as resource availability, predation pressure, and inter- and intraspecific competition [43]. During dispersal, distance-dependent mortality occurs, i.e., the further two patches are apart, the more biomass is lost to the hostile matrix separating them. We constructed 30 model food webs and simulated each food web on 72 different landscapes. For each simulation we generated landscapes on two independent gradients covering two aspects of fragmentation, namely number of patches and habitat isolation (figure 1c). We achieved a full range for the gradient of habitat isolation (landscape connectance ranging from 0 to 1, figure 3c). Additionally, we performed dedicated simulation runs to reference the two extreme cases, i.e. (1) landscapes in which all patches are direct neighbours without a hostile matrix, and thus, no dispersal mortality, and (2) fully isolated landscapes, in which no species can bridge between patches, and thus, a dispersal mortality of 100%. Additionally, we tested a null model in which all species have the same maximum dispersal distance. To visualise the impact of number of patches and habitat isolation on species diversity we used GAMMs from the mgcv package in R [44]. See the supplement for detailed information on the maximum dispersal distance, the additional simulations and the statistical analysis.

## Results

### Species diversity patterns

Our simulation results identify habitat isolation (defined as the mean distance between habitat patches, 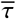, figure 2, x-axis) as the key factor driving species diversity loss. As expected, we find fewer species on patches (the averaged local diversity, 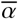) in landscapes in which habitats are highly isolated (figure 2, left panel). In contrast to the decrease in 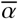-diversity, *β*-diversity (figure 2, middle panel), which describes differences in the community composition between patches, increases with habitat isolation. This increase happens around the infliction point of the landscape connectance at a mean patch distance log_10_ 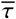 ≈ of −0.5, at which 50% of all possible patch to patch connections are lost (supplement figure S4, first panel). *γ*-diversity, the species diversity in the landscape, shows a more complicated pattern. First it decreases due to the loss of 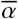-diversity with habitat isolation. This decrease is then reversed by the increase of *β*-diversity and the *γ*-diversity increases again with habitat isolation (figure 2, right panel). The number of habitat patches in a landscape, *Z* (figure 2, y-axis), only marginally affects the diversity patterns. The additional simulations of the two extreme cases (i.e. joint scenario with no dispersal loss and fully isolated scenario with 100% dispersal mortality) support these patterns (see the supplement, section S7 for the corresponding results). We further show that the isolation-induced species loss also translates into a loss of trophic complexity, i.e. isolated landscapes are characterised by reduced food webs with fewer species and fewer trophic levels (see the supplement, figure S2).

**Figure 2:**
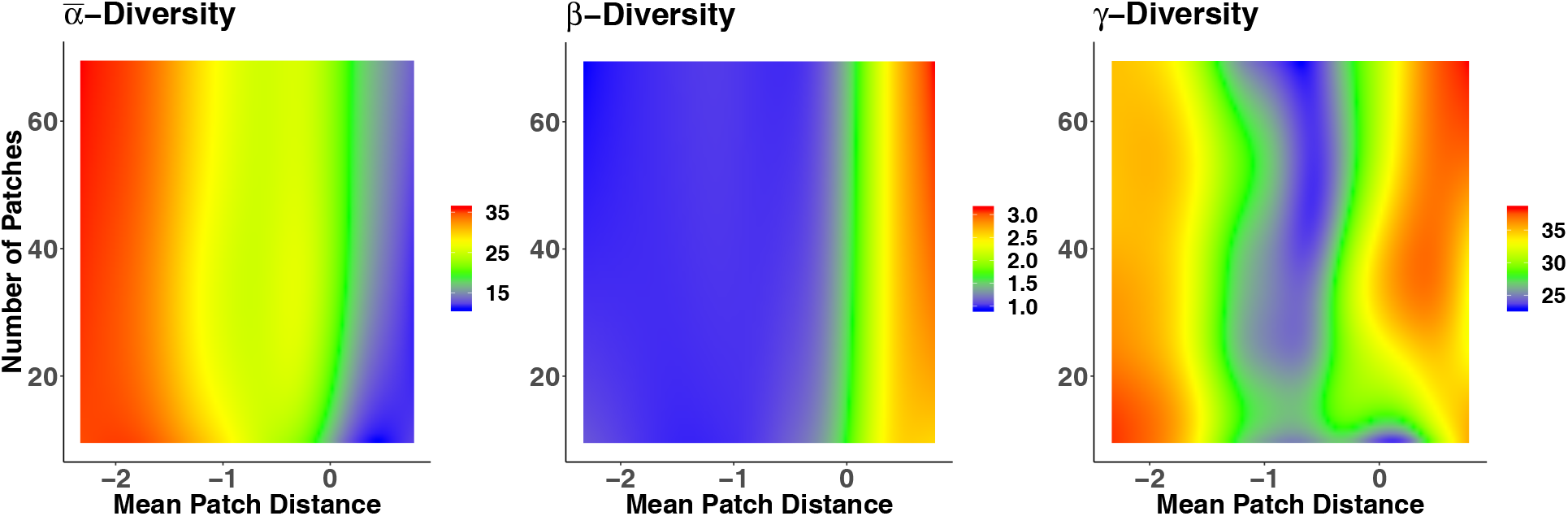
Heatmaps visualising 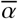-, *β*- and *γ*-diversity (colour-coded; z-axis) in response to habitat isolation, i.e. the mean patch distance (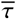, log_10_-transformed; x-axis) and the number of habitat patches (*Z*; y-axis), respectively. We generated the heatmaps based on the statistical model predictions (see the methods section).

### Differences among trophic levels

As the number of patches only marginally affects species diversity patterns, we hereafter focus on the effects of habitat isolation on trophic-dependent differences among species (figure 3). In figure 3, biomass densities, *B*_i_, and landscape connectances, *ρ*_*i*_, represent the average of each species *i* over all food webs. Species are ranked according to their body mass. Thus, although species body masses differ between food webs, species 1 is always the smallest, species 2 the second smallest and so forth. The same applies to *ρ*_*i*_, where the landscape connectance of consumer species is body mass dependent, but the connectance of plant species is body mass independent (see the methods section). In well-connected landscapes (i.e. landscapes with small mean patch distances, 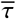), large and medium-sized consumer species (except the very largest) have higher population biomass densities than smaller consumers (figure 3a,c). With expanding distances between habitat patches, large-bodied consumers at high trophic positions (figure 3a, red to blue lines) show a particularly strong decrease in population biomass densities. Small consumer species (figure 3a, orange to yellow lines) are generally less affected by increasing habitat isolation. Plant species show a less consistent response to increased isolation, with most species slightly increasing their biomass density (figure 3b, green lines). Based on our assumption that the maximum dispersal distance of animals scales with body mass, the ability to connect a landscape follows the same allometric scaling (figure 3c). Despite this dispersal advantage, intermediate-sized and large animal species (figure 3a, red to blue lines) lose biomass in landscapes in which they still have the potential to fully connect (almost) all habitat patches (figure 3c). The differences in plant species biomass densities cannot be attributed to body mass dependent species-specific dispersal distances as for plants maximum dispersal distances were randomly assigned, and thus, there is no connection between body mass and landscape connectance (*ρ*_*i*_, figure 3d). Additional simulations, in which we assumed a constant maximum dispersal distance for all species of *δ*_*i*_ = *δ*_*max*_ = 0.5, support the negligibility of species-specific differences in dispersal ability for the emerging diversity patterns (see the supplement, figure S3).

**Figure 3:**
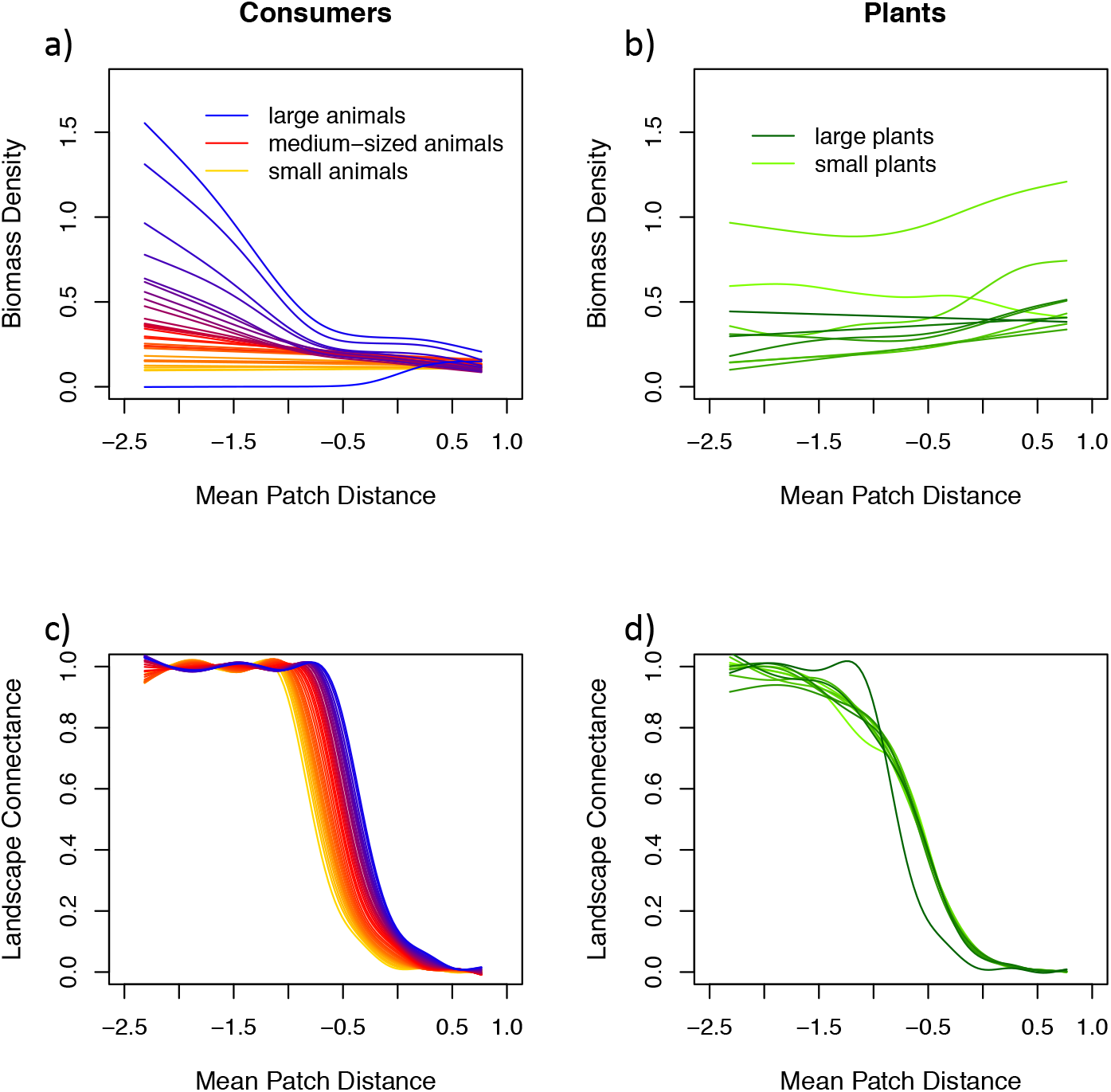
Top row: Mean biomass densities [log10(biomass density-1)] of animal consumer species and basal plant species (b) over all food webs (*B*_i_, log_10_-transformed; y-axis) in response to habitat isolation, i.e. the mean patch distance (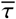, log_10_-transformed; x-axis). Each colour depicts the biomass density of species *i* averaged over all food webs: (a) colour gradient where orange represents the smallest, red the intermediate and blue the largest consumer species; (b) colour gradient where light green represents the smallest and dark green the largest plant species. Bottom row: Mean species-specific landscape connectance (*ρ*_*i*_; y-axis) for consumer (c) and plant species (d) over all food webs as a function of the mean patch distance (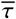, log_10_-transformed; x-axis). See the supplement figure S9 for standard errors in biomass densities for four exemplary species.

## Discussion

Habitat fragmentation is a major driver of global biodiversity decline. To date, a comprehensive understanding of how the different aspects of habitat fragmentation, i.e. habitat loss [6], number of fragments and isolation, affect the diversity patterns of species embedded in complex ecological networks such as food webs is lacking (see e.g. meta-analysis by Martinson and Fagan [15], and references therein). Our simulation experiment allows us to independently explore the effects of number of fragments (i.e., number of habitat patches in the landscape), and of habitat isolation (i.e., distance between patches) on persistence and biomass densities of species in complex communities. We identified habitat isolation to be responsible for species diversity decline both at the local and regional scale. The rate at which a species loses biomass density strongly depends on its trophic position. Large-bodied consumer species at the top of the food web are most sensitive to isolation although they are dispersing most effectively (i.e. for them, increasing distances between habitat patches do not necessarily result in the loss of dispersal pathways or a substantial increase of dispersal mortality). Surprisingly, we find top species to loose biomass density and sometimes even go extinct in landscapes they can still fully connect, whereas the biomass densities of small consumer species at lower trophic levels and plant species are only marginally affected by increasing habitat isolation. We attribute the accelerated loss of top species to the energy limitation propagated through the food web: with increasing habitat isolation an increasing fraction of the biomass production of the lower trophic levels is lost due to mortality during dispersal and is thus no longer available to support the higher trophic levels. Additionally, the reduced top-down pressure on smaller consumers seems to compensate for their increased dispersal loss. Our model adds a complementary perspective to previous research pointing towards a trophic-dependent extinction risk due to constraints in resource availability with increasing habitat fragmentation [9, 38].

### Habitat isolation drives species loss

The increasing isolation of habitat fragments poses a severe threat to species persistence (but see [45, 46]). We demonstrate in our simulation experiment that the generally observed pattern of species loss with increasing habitat isolation (e.g. [3]) also holds for species embedded in large food webs. The loss of species occurs both at the local (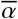-diversity) and regional (*γ*-diversity) scale. For the latter, however, an increase in *β*-diversity compensates the loss in local diversity 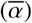 when landscapes become very isolated and *γ*-diversity increases again (see below, Habitat isolation promotes *β*-diversity).

We modelled dispersal between habitat patches by assuming an energy loss for the dispersing organisms – a biologically realistic assumption as landscape degeneration, which often occurs concurrently with habitat fragmentation, increases the hostility of the habitat matrix [3]. Consequently, the dispersal mortality, and thus, biomass loss of populations to the habitat matrix increases substantially when dispersal distances between habitat patches expand. To account for the variation in dispersal ability among trophic groups, we incorporated species-specific maximum dispersal distances. For animal species, this maximum dispersal distance increases like a power law with body mass, therefore weakening the direct effect of habitat isolation the larger a species is. Despite this, top predators and other large consumer species respond strongly to isolation. These species exhibit a dramatic loss in biomass density or even go extinct in landscapes they still perceive as almost fully connected (landscape connectance, *ρ*_*i*_, close to one), which indicates that their response to habitat isolation is mediated by indirect effects originating from the local food web dynamics.

### Local food web dynamics and energy limitation drive top predator loss

In local food webs energy is transported rather inefficiently from the basal to the top species, with transfer efficiency in natural systems often only around 10% [47]. This energy limitation effectively controls the food chain length [26] and renders large species at high trophic levels vulnerable to extinction due to resource shortage [48]. In our model, energy availability decreases if habitat isolation is high as this increases biomass loss during dispersal. This affects particularly small species at lower trophic levels since they generally have the highest metabolic costs per unit biomass and therefore the highest biomass losses per distance travelled [33, 41]. The biomass loss during dispersal consequently reduces the net biomass production at the bottom of the food web and severely threatens species at higher trophic positions that already operate on a very limited resource supply.

Moreover, due to the feedback mechanisms regulating the community dynamics within complex food webs, a loss of top consumer species can have severe consequences for the functioning and stability of the network [21, 22]. A loss of top-down regulation can, for instance, lead to secondary extinctions resulting in simpler food webs [21, 49] – an additional mechanism that can foster the loss of biodiversity as observed in our simulations. However, we also see a much more direct effect of the changing community composition: The biomass densities of small species that suffer most from increased dispersal mortality do not, as one might expect, decline much as isolation progresses. We attribute this to a release from top-down control as their consumers lose biomass or even go extinct, which counters the negative direct effect of habitat isolation. These arguments suggest that differential dispersal capabilities are less important than energetic limitations in explaining the strong negative response of large consumers to habitat isolation. This claim is supported by the additional simulations where all species experienced the same level of dispersal mortality, which yielded similar results (see the supplement, figure S3).

We did not find an effect of the number of patches on 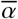-, *β*- and *γ*-diversity. As we model biomass densities on patches without defined area (see below, Model specifications), fewer patches do not reflect habitat loss, but rather the loss of fragments, i.e. stepping stones in the dispersal network. Thus the energy limitation in our simulated landscapes derives from direct dispersal loss and cascading effects of dispersal losses of resources. For plant and small animal species this can be understood easily, as these species are less energy limited and thus are able to persist on a single habitat patch. For larger animal species the situation is more subtle: While they can integrate over multiple patches, feeding interactions still always occur on one patch at a time. If the biomass densities of their resources (and thus also the realised feeding rate) is too low on a particular patch to cover their metabolic requirements, they gain no advantage from the addition of more patches with equally low resource abundance.

### Habitat isolation promotes *β*-diversity

Contrary to the decline in 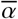-diversity with increasing habitat isolation, we find an increase in *β*-diversity starting from around log_10_ mean patch distance 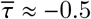. We assumed identical abiotic conditions on all habitat patches, i.e. there are no differences in nutrient availability or background mortality rates. Therefore, any differences in conditions experienced by the species on different patches can only originate from the initial community composition and the structure of the dispersal network. One way for such different conditions to emerge is the disintegration of the dispersal network into several smaller clusters. Up to a log_10_ mean patch distance 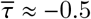, the species with the largest maximum dispersal distance (which could be both large animals that have not already gone extinct and plants with a randomly selected large dispersal distance) have a landscape connectance (*ρ*_*i*_) of at least 0.5. This dispersal advantage easily allows them to connect all patches to a single network component, thereby providing homogenisation for the meta-food-web. However, as the mean patch distance increases further, even these species cannot bridge all gaps in the habitat matrix any more and clusters of patches emerge that are for all species disconnected from the other patches. As these clusters vary in the number of patches and mean patch distance within the cluster, the level of dispersal mortality experienced by the species on the different clusters can also vary considerably. Any further increase in mean patch distance causes the landscape connectance to drop to nearly zero for all species and all patches within the landscape approach complete isolation. With no immigration into isolated patches, non-resident species cannot colonise them and initial community compositions drive dissimilarities among patches. However, the initial *β*-diversity is not sufficient in explaining the high *β*-diversity in strongly isolated landscapes (supplement figure S4). This suggests that different food web positions of initial species lead to different cascading effects in local food web dynamics with more or less secondary extinctions on isolated patches further increasing differences in local community compositions. The increase in *β*-diversity is even stronger than the loss of local diversity resulting in an increase in *γ*-diversity in highly isolated landscapes. However, species contributing to this high *γ*-diversity tend to occur on fewer patches and thus are more prone to go extinct in the whole landscape due to stochastic extinction events.

### Model specifications

The framework we propose here for modelling meta-food-webs is very general and allows for a straightforward implementation of future empirical insight where we so far had to rely on plausible assumptions. The trophic network model for the local food webs is based on a tested and realistic allometric framework [41] with a fixed number of 40 species – a typical value in dynamic food web modelling (e.g. [50, 51]). We based all model parameters on allometric principles [33, 52] allowing for a simple adaptation of our modelling approach to other trophic networks such as empirically sampled food webs [53] or other food web models such as the niche model [54]. Moreover, empirical patch networks (e.g. the coordinates of meadows in a forest landscape) or other dispersal mechanisms [6, 55] may be incorporated in the future. In our simulations, biomass loss during dispersal is predominantly responsible for the decline in species diversity. We linked the maximum dispersal distance of animals and thereby also their mortality during dispersal to body mass, which is plausible because larger animal species can move faster [34], and thus, have to spend less time in the hostile habitat matrix. Interestingly, however, we did not find any empirical study relating body mass directly to mortality or biomass loss during migration. If such information becomes available in the future, it can be easily incorporated into our modelling framework. Further, we deliberately assumed all habitat patches to share the same abiotic conditions [56] as we wanted to focus on the general effects of the interaction of complex food web and dispersal dynamics. Adding habitat heterogeneity among patches, e.g. by modifying nutrient availability or mean temperature, however, is straightforward and can be expected to yield additional insight into the mechanisms for the maintenance of species diversity in meta-food-webs. Finally, by using a dynamics model formulated in terms of biomass densities instead of absolute biomasses (or population sizes), we make the implicit assumption that patches do not have an absolute size. Thus, the number of patches in a landscape cannot be directly linked to the total amount of habitat but rather reflects the number of fragments, i.e. stepping stones in the dispersal network of a species. A decreasing number of patches thus does not necessarily imply habitat loss. In order to also address effects of habitat loss (in terms of area), the model could be adapted to include for example area specific extinction thresholds and absolute biomasses in dispersal dynamics, but this was beyond the scope of this study.

### Synthesis and outlook

Our simulation experiment demonstrates that habitat isolation reduces species diversity in complex food webs in general, with differences in the effect across trophic levels. In increasingly isolated landscapes, energy becomes limited, which decreases the biomass density of large consumers or even drives them extinct. These primary extinctions may result in a cascade of secondary extinctions, given the importance of top predators for food web stability [24, 57]. The increased risk of network downsizing, i.e. simple food webs with fewer and smaller species [14, 58], stresses the importance to consider both direct and indirect trophic interactions as well as dispersal when assessing the extinction risk of species embedded in complex food webs and other ecological networks.

To date, most conservation research focuses on single species and does not consider the complex networks of interactions in natural communities [7, 14]. However, the patterns we presented here clearly support previous studies highlighting the importance of trophic interactions (e.g. [9, 37, 38]). We show that the fragmentation-induced extinction risk of species strongly depends on their trophic position, with top species being particularly vulnerable. Given that top-down regulation can stabilise food webs [24, 57], the loss of top predators might entail unpredictable consequences for adjacent trophic levels, destabilise food webs, reduce species diversity and trophic complexity and ultimately compromise ecosystem functioning [23, 24]. In addition to the trophic position of a species, the trophic structure of the food web has also been shown to be an important aspect (see [11]). Our results suggest that bottom-up energy limitation caused by dispersal mortality due to habitat isolation can be a critical factor driving species loss and the reduction of trophic complexity. The extent of this loss strongly depends on the spatial context (see also [6]). Thus, to maintain species-rich and trophic-complex natural communities under future environmental change, effective conservation planning must consider this interdependence of spatial and trophic dynamics. Notably, conservation planning should also consider habitat isolation and matrix hostility (and consequently dispersal mortality) to ensure sufficient biomass exchange between local populations, capable of inducing spatial rescue effects, and to alleviate bottom-up energy limitation of large consumers. Energy limitations can also result from habitat loss (which we did not model here), decreasing energy availability at the bottom of the food web affecting local dynamics intrinsically independent of dispersal. Thus, avoiding habitat loss remains a crucial aspect [2, 46]. We highlight the need to explore food webs and other complex ecological networks in a spatial context to achieve a more holistic understanding of biodiversity and ecosystem processes.

## Supporting information

Supplement Material - Methods and sensetivity analysis

## Acknowledgements

This study was financed by the German Research Foundation (DFG) in the framework of the research unit FOR 1748 - Network on Networks: The interplay of structure and dynamics in spatial ecological networks (RA 2339/2-2, BR 2315/16-2, GU 1645/1-1). Further, JH, RR, UB and BCR gratefully acknowledge the support of the German Centre for Integrative Biodiversity Research (iDiv) Halle-Jena-Leipzig funded by the German Research Foundation (FZT 118). The scientific results have (in part) been computed at the High-Performance Computing Cluster EVE of the Helmholtz Centre for Environmental Research - UFZ and iDiv, and we thank the staff of EVE (in particular Christian Krause from iDiv) for their support. Furthermore, we thank Thomas Boy for his technical support and assistance with programming issues.

## Author contributions

All authors conceived and designed the modelling framework; JH and RR ran the simulations on the high-performance-cluster; RR analysed the data with support from all other authors; all authors contributed to interpreting the results; JH wrote the first draft of the manuscript with support from RR and MS and JH and RR led the editing. All authors contributed critically to the drafts and gave final approval for publication.

## Data accessibility

We enable full reproducibility of our study by providing the original C- and R-code on Dryad. https://datadryad.org/review?doi=doi:10.5061/d

## Competing interests

The authors declare no competing interests.

## Notes

#### Summary of Updates

Rerun of simulations which clarified the results. This was changed in section results accordingly; Reduce the method part to a method summary. An elaborate method description is provided in the supplementary material; Clarify the relatedness to fragmentation and use of coherent wording which is used in the literature in this context.

