## Supplement Material - Methods and sensetivity analysis for "The biggest losers: Habitat isolation deconstructs complex food webs from top to bottom"

### S1 Food web and local population dynamics

We consider a multitrophic metacommunity consisting of 40 species on a varying number of randomly positioned habitat patches, $Z$ (the meta-food-web, figure 1b). All patches have the same abiotic conditions and each patch can potentially harbour the full food web, consisting of 10 basal plant and 30 animal consumer species. The feeding links (i.e. who eats whom) are constant over all patches (figure 1a,b) and are as well as the feeding dynamics determined by the allometric food web model by Schneider *et al.* [1]. We integrate dispersal as species-specific biomass flow between habitat patches (figure 1b,d).

Using ordinary differential equations to describe the feeding and dispersal dynamics, the rate of change in biomass density, $B_{i,z}$, of species $i$ on patch $z$ is given by

$$\frac{dB_{i,z}}{dt} = T_{i,z} - E_{i,z} + I_{i,z}\,, \tag{1}$$

with $T_{i,z} = \upsilon_{i,z} \cdot B_{i,z}$ as the rate of change in biomass density determined by local feeding interactions (where $\upsilon_{i,z}$ is the per capita growth rate, see table S2), $E_{i,z}$ as the total emigration rate of species $i$ from patch $z$ (equation 2), and $I_{i,z}$ as the total rate of immigration of species $i$ into patch $z$ (equation 4).

#### Local food web dynamics

We use an allometric trophic network model (ATN model) based on the work of Schneider *et al.* [1] & Kalinkat *et al.* [2] to simulate the trophic dynamics of local populations ($T_{i,z}$ in Equation 1). Regarding this term, we distinguish between animal species (Equation T1-1) and basal plant species (Equation T1-6). In each patch, the biomass dynamics of animal species (biomass densities $A_{i,z}$) is given by the differences between growth due to consumption of animal or plant species and losses due to mortality through predation and metabolic demands. The rate of change in plant biomass densities $P_{i,z}$ depends on the uptake of the two resources, mortality through grazing, and also accounts for metabolic losses. We used a dynamic nutrient model (equation T1-8) with two nutrients (concentrations $N_{l,z}$) of different importance as the energetic basis of our food web [1, 3].

The topological network model is an extension of the niche model originally introduced by Williams & Martinez [4] and accounts for allometric degree distributions and recent data on scaling relationships for species body mass and trophic levels [5]. Each species $i$ is fully characterised by its average adult body mass $m_i$. We sampled $\log_{10}$ body masses of animal species randomly with a uniform probability density from the inclusive interval $(2, 12)$ and the $\log_{10}$

body masses of plant species from the inclusive interval $(0, 6)$ (for empirical examples see [6]). This step makes the model inherently stochastic, but from hereon, all other steps are completely deterministic. The model is designed such that animal consumers feed on resources, which can be both plants and other animal species that are smaller than themselves. Body masses further determine the interaction strengths of feeding links as well as the metabolic demands of species.

Data from empirical feeding interactions are used to parametrise the functions that characterise the optimal prey body mass and the location and width of the feeding niche of a predator. From each $m_i$ a unimodal attack kernel, called feeding efficiency, $L_{ij}$, is constructed which determines the probability of consumer species $i$ to attack and capture an encountered resource species $j$. We model $L_{ij}$ as an asymmetrical hump-shaped Ricker's function (equation T1-4) that is maximised for an energetically optimal resource body mass (optimal consumer-resource body mass ratio $R_{opt} = 100$) and has a width of $\gamma = 2$. The maximum of the feeding efficiency $L_{ij}$ equals 1. Table S1 list the full set of equation and table S2 is an overview of the standard parameter set for the equations. See also Schneider *et al.* [1] for further information regarding the allometric food web model.

Table S1: Ordinary differential equations describing the local population dynamics driven by feeding interactions (see Schneider *et al.* [1]. We use the same allometric constraints and parameter ranges.

| Equation No. | Model equations | Description |
|---|---|---|
| Equation T1-1 | Animal population dynamics $$\frac{dA_{i,z}}{dt} = e_P A_{i,z} \sum_j F_{ij,z} + e_A A_{i,z} \sum_k F_{ik,z} - \sum_k A_{k,z} F_{ki,z} - x_i A_{i,z}$$ | Rate of change of biomass density of animal species $i$ on patch $z$; with conversion efficiency $e_P = 0.545$ typical for herbivory [7]; conversion efficiency $e_A = 0.906$ typical for carnivory [7]; feeding rate $F_{ij,z}$ of consumer $i$ on resource $j$ on patch $z$; metabolic demands per unit biomass for animals $x_i = x_A m_i^{-0.305}$ with scaling constant $x_A = 0.314$ [8, 9]. The first sum goes over all plant resources $j$, the second over all animal resources $k$ and the third over all animal predators $k$ of animal species $i$. |
| Equation T1-2 | Functional response $$F_{ij,z} = \frac{\omega_i b_{i,j} R_{j,z}^{1+q}}{1 + cA_{i,z} + \omega_i \sum_k b_{ik} h_{ik} R_{k,z}^{1+q}} \cdot \frac{1}{m_i}$$ | Per unit biomass feeding rate of consumer $i$ as function of its own biomass density, $A_i$, (taking interference competition $c$, which is the time lost due to intraspecific encounters, sampled from a normal distribution with mean $\mu_c = 0.8$ and s.d. $\sigma_c = 0.2$ for each food web), and biomass density of the resource $R_j$ (either animal $A_j$ or plant species $P_j$); with $b_{ij}$, resource specific capture coefficient (Eq. T1-3); $h_{ij}$, resource-specific handling time (Eq. T1-5); $\omega_i = 1/(\text{number of resource species of } i)$, relative consumption rate accounting for the fact that a consumer has to split its consumption if it has more than one resource species. |

Continued on next page

**Table S1 – continued from previous page**

| Equation No. | Model equations | Description |
|---|---|---|
| Equation T1-3 | Capture coefficient $b_{ij} = a_k m_i^{\beta_i} m_j^{\beta_j} L_{ij}$ | Resource specific capture coefficient of consumer species $i$ on resource species $j$ scaling the feeding kernel $L_{ij}$ by a power function of consumer and resource body mass, assuming that the encounter rate between consumer and resource scales with their respective movement speed. We sample the exponents $\beta_i$ and $\beta_j$ from normal distributions (mean $\mu_{\beta_i} = 0.42$, s.d. $\sigma_{\beta_i} = 0.05$; $\mu_{\beta_j} = 0.19$, s.d. $\sigma_{\beta_j} = 0.04$, respectively [10]). We divide here the group of consumer species into the subgroup of carnivorous and herbivorous species each comprising a constant scaling factor for their capture coefficients $a_k$ with $k \in 0, 1$ ($a_0 = 40$ for carnivorous species and $a_1 = 5000$ for herbivorous species); For plant resources, $m_j^{\beta_j}$ was replaced with the constant value of 1 (as plants do not move). |
| Equation T1-4 | Feeding efficiency $L_{ij} = \left(\frac{m_i}{m_j R_{opt}} e^{1-\frac{m_i}{m_j R_{opt}}}\right)^{\gamma}$ | The probability of consumer $i$ to attack and capture an encountered resource $j$ (which can be either plant or animal), described by an asymmetrical hump-shaped curve (Ricker's function), with width $\gamma$ = 2 centered around an optimal consumer-resource body mass ratio $R_{opt} = 100$. |
| Equation T1-5 | Handling time $h_{ij} = h_0 m_i^{\eta_i} m_j^{\eta_j}$ | The time consumer $i$ needs to kill, ingest and digest resource species $j$, with scaling constant $h_0 = 0.4$ and allometric exponents $\eta_i$ and $\eta_j$ drawn from normal distributions with means $\mu_{\eta_i} = -0.48$ and $\mu_{\eta_j} = -0.66$, and standard deviations $\sigma_{\eta_i} = 0.03$ and $\sigma_{\eta_i} = 0.02$, respectively [11]. |

Continued on next page

**Table S1 – continued from previous page**

| Equation No. | Model equations | Description |
|---|---|---|
| Equation T1-6 | Plant population dynamics $\frac{dP_{i,z}}{dt} = r_i G_i P_{i,z} - \sum_{k} A_{k,z} F_{ki,z} - x_i P_{i,z}$ | Rate of change of biomass density of plant species $i$ on patch $z$; with predation loss $F_{ki,z}$ summed over all consumer species $k$ feeding on plant species $i$; metabolic demands per unit biomass for plants $x_i = x_P m_i^{-0.25}$ with $x_P = 0.138$; intrinsic growth rate $r_i = m_i^{-0.25}$; species specific growth factor $G_i$ (Eq. T1-7). |
| Equation T1-7 | Growth factor for plants $G_i = min\left(\frac{N_1}{K_{i,1} + N_1}, \frac{N_2}{K_{i,2} + N_2}\right)$ | Species-specific growth factor of plants determined dynamically by the most limiting nutrient $l \in 1, 2$; with $K_{i,l}$, half-saturation densities determining the nutrient uptake efficiency assigned randomly for each plant species $i$ and nutrient $l$ (uniform distribution within $(0.1, 0.2)$). The term in the minimum operator approaches 1 for high nutrient concentrations. |
| Equation T1-8 | Nutrient dynamics $\frac{dN_{l,z}}{dt} = D(S_l - N_l) - \nu_l \sum_{i,z} r_i G_i P_{i,z}$ | Rate of change of nutrient concentration $N_l$ of nutrient $l \in \{1, 2\}$ on patch $z$, with global turnover rate $D = 0.25$, determining the rate at which nutrients are refreshed; supply concentration $S_l$, determining the maximum nutrient level of each nutrient, $l$, drawn from normal distributions with mean $\mu_S = 50$ and standard deviation $\sigma_S = 2$ (provided $S_l > 0$); relative nutrient content in plant species biomass $\nu_l$ ($\nu_1 = 1$, $\nu_2 = 0.5$). |

### S2 Generating landscapes

We generated differently fragmented landscapes, represented by random geometric graphs [12], by randomly drawing the locations of $Z$ patches from a uniform distribution between 0 and 1 for x- and y-coordinates respectively. We created landscapes of different size by scaling the maximum dispersal distance of all organisms $\delta_{max}$ with a factor, $Q$, to represent landscape sizes with edge lengths between 0.01 and 10. We obtained the number of patches, $Z$, by using a stratified random sampling approach, i.e. we added a random number drawn from an integer uniform distribution between 0 and 9 to a series of numbers of $10, 20, \ldots, 60$. Similarly, we set the landscape size, $Q$, by adding a random number drawn from a uniform distribution between 0 and 1 (respectively 0 and 0.1 for landscape sizes below 1) to a series of numbers of $0.01, 0.1, 0.2, 0.3, 0.5, 0.7, 0.9, 1, 3, 5, 7, 9$.

### S3 Dispersal

We model dispersal between local communities as a dynamic process of emigration and immigration, assuming dispersal to occur at the same timescale as the local population dynamics [13]. Thus, biomass flows dynamically between local populations and the dispersal dynamics directly influence local population dynamics and vice versa [14]. Similar approaches have been used by e.g. Abrams & Ruokolainen [15] and Ims & Andreassen [16]. We model a hostile matrix between habitat patches that does not allow for feeding interactions to occur during dispersal, and thus, assume the biomass lost to the matrix to scale linearly with the distance travelled.

**Emigration** The total rate of emigration of species $i$ from patch $z$ is

$$E_{i,z} = d_{i,z} B_{i,z} \,, \tag{2}$$

with $d_{i,z}$ as the corresponding per capita dispersal rate. We model $d_{i,z}$ as

$$d_{i,z} = \frac{a}{1 + e^{b(x_i - \upsilon_{i,z})}} \,, \tag{3}$$

with $a$, the maximum dispersal rate, $b$, a parameter determining the shape of the dispersal rate (figure S1), $x_i$, the inflection point determined by the metabolic demands per unit biomass of species $i$, and $\upsilon_{i,z}$, the per capita net growth rate of species $i$ on patch $z$. We chose to model $d_{i,z}$ as a function of each species' per capita net growth rate to account for emigration triggers such as resource availability, predation pressure and inter- and intraspecific competition [14, 17]. If for example an animal species' net growth is positive, there is no need for dispersal and emigration will be low. However, if the local environmental conditions deteriorate, the growing incentives to search for a better habitat increase the fraction of individuals emigrating. For plants, we assumed an additional scenario as there are examples of different life history strategies. There are for example plant species which disperse from their local habitat when they are doing well, i.e. they have a high net growth rate, as they can allocate more resources into reproduction resulting in higher seed dispersal [18]. However, there are also examples where plants reallocate resources into reproduction when they are doing poorly [19] (figure S1b).

For each simulation run, $a$ was sampled from a Gaussian distribution ($\mu_{aS}, \sigma_{aS}$) and $b$ was sampled from an integer uniform distribution within inclusive limits that differed between consumer and plant species (see table S2). The different intervals reflect different dispersal triggers for animals and plants.

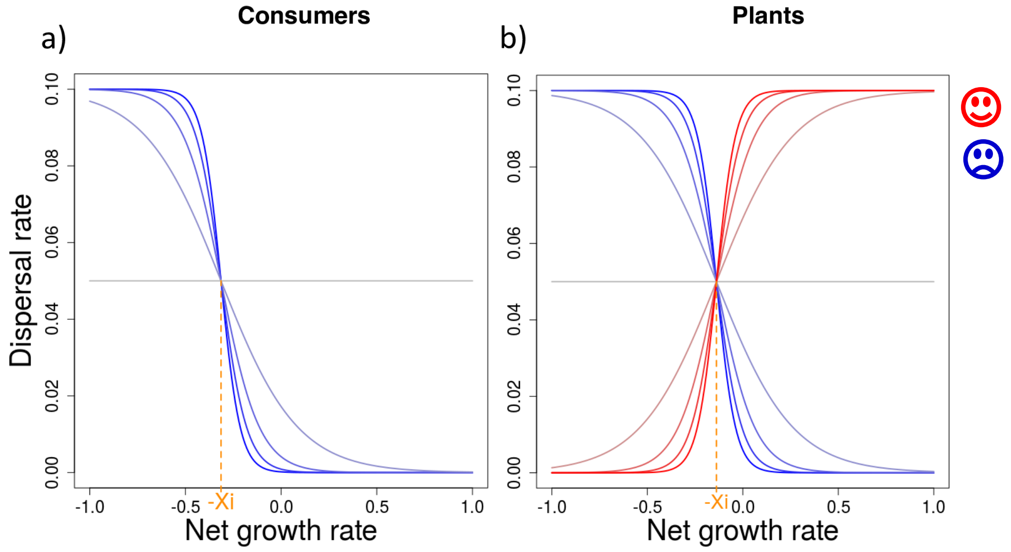


Figure S1: Functions illustrating the dispersal rate $d_i$ for animal (a) and plant species (b), where $x_i$ marks the inflection point for each species $i$ determined by the metabolic demands per unit biomass of species $i$ (see Table S1). The colours blue and red represent different dispersal strategies and the respective colour gradients depict the parameter range of $b$, which determines the slope of the dispersal rate (see equation 3 in the manuscript). For the purpose of illustration, we set the maximum dispersal rate to $a = 0.1$ and for animals and plants $x_{i_A} = 0.314$ and $x_{i_P} = 0.1384$, respectively.

**Immigration** The rate of immigration of biomass density of species $i$ into patch $z$ follows

$$I_{i,z} = \sum_{n \in N_z} E_{i,n}(1 - \delta_{i,nz}) \frac{1 - \delta_{i,nz}}{\sum_{m \in N_n} 1 - \delta_{i,nm}}, \tag{4}$$

where $N_z$ and $N_n$ are the sets of all patches within the dispersal range of species $i$ on patches $z$ and $n$, respectively. In this equation, $E_{i,n}$ is the emigration rate of species $i$ from patch $n$, $(1 - \delta_{i,nz})$ is the fraction of successfully dispersing biomass, i.e. the fraction of biomass not lost to the matrix, and $\delta_{i,nz}$ is the distance between patches $n$ and $z$ relative to species $i$'s maximum dispersal distance $\delta_i$ (see below paragraph Maximum dispersal distance). The term $\frac{1-\delta_{i,nz}}{\sum 1-\delta_{i,nm}}$ determines the fraction of biomass of species $i$ emigrating from source patch $n$ towards target patch $z$. This fraction depends on the relative distance between the patches, $\delta_{i,nz}$, and the relative distances to all other potential target patches $m$ of species $i$ on the source patch $n$, $\delta_{i,nm}$. Thus, the flow of biomass is greatest between patches with small distances. For numerical reasons, we did not allow for dispersal flows with $I_{i,z} < 10^{-10}$. In this case, we immediately set $I_{i,z}$ to 0.

**Maximum dispersal distance** Based on empirical observations (e.g. [20]) and previous theoretical frameworks (e.g. [10, 21–23]), we assume that the maximum dispersal distance $\delta_i$ of animal species increases with their body mass. For animal species, the body mass $m_i$ determines how fast and how far they can travel through the matrix before needing to rest and feed in a habitat patch. Thus animal species at high trophic positions can disperse further than smaller animals at lower trophic levels. Each animal species perceives its own dispersal network dependent on its species-specific maximum dispersal distance

$$\delta_i = \delta_0 m_i^{\epsilon}\,, \tag{5}$$

where the exponent $\epsilon = 0.05$ determines the slope of the body mass scaling of $\delta_i$. We chose a positive value for $\epsilon$ to account for a higher mobility of animals with larger body masses. The intercept $\delta_0 = 0.1256$ was chosen such that the animal species with the largest possible body mass of $m_i = 10^{12}$ had a maximum dispersal distance of $\delta_i = 0.5$. Thus, the animal species with the smallest possible body mass of $m_i = 10^2$ had a maximum dispersal distance of $\delta_i = 0.158$.

As plants are passive dispersers driven by e.g. wind with no clear relationship between body mass and dispersal distance, we model their maximum dispersal distance as random and body mass independent [20]. We sampled $\delta_i$ for each plant species from a uniform probability density within the interval $(0, 0.5)$. Thus, the best plant disperser can potentially have the same maximum dispersal distance as the largest possible animal species (table S2). Additionally, we tested a null model in which all species have the same maximum dispersal distance of $\delta_i = \delta_{max}$. See section S8 for further information on the additional simulations.

Table S2: Model parameters and output variables.

| Parameter | Description | Value |
|---|---|---|
| | **Trophic interactions between species** | |
| $e_A$ | conversion efficiency animal species | 0.906; [7] |
| $e_P$ | conversion efficiency plant species | 0.545; [7] |
| $x_A$ | scaling constant metabolic demands animal species | 0.314; [9] |
| $x_P$ | scaling constant metabolic demands plant species | 0.138; [9] |
| $\mu_c$, $\sigma_c$ | mean and standard deviation for interference competition | 0.8, 0.2 |
| $a_0$ | scaling factor capture coefficient for carnivorous species | 40 |
| $a_1$ | scaling factor capture coefficient for herbivorous species | 5000 |
| $\mu_{\beta_i}$, $\sigma_{\beta_i}$ | mean and standard deviation allometric exponent for attack rates consumer | 0.42, 0.05; [10] |
| $\omega_i$ | relative consumption rate | $\frac{1}{\text{number of prey species } i}$ |
| $R_{opt}$ | optimal consumer-resource body mass ratio | 100 |
| $\gamma$ | scaling exponent Ricker's function | 2 |
| $h_0$ | scaling factor handling time | 0.4 |
| $\mu_{\eta_i}$, $\sigma_{\eta_i}$ | mean and standard deviation allometric exponent handling time consumer | -0.48, 0.03; [11] |
| $\mu_{\eta_j}$, $\sigma_{\eta_j}$ | mean and standard deviation allometric exponent handling time resource | -0.66, 0.02; [11] |
| $\mu_q$, $\sigma_q$ | mean and standard deviation hill coefficient | 1.5, 0.2 |
| | **Nutrient dynamics** | |
| $K$ | half saturation density nutrient uptake | (0.1, 0.2) |
| $D$ | nutrient turnover rate | 0.25 |
| $\mu_{S_l}$, $\sigma_{S_l}$ | mean and standard deviation of nutrient supply concentration | 50, 2 |
| $v_1$, $v_2$ | relative nutrient content in plant species biomass | 1, 0.5 |
| | **Dispersal dynamics** | |
| $\delta_{max}$ | species-specific maximum dispersal distance | 0.5 |
| $\epsilon$ | scaling exponent for species-specific maximum dispersal distance | 0.05 |
| $\mu_{a_S}$, $\sigma_{a_S}$ | mean and standard deviation of max. emigration | 0.1, 0.03 |
| $\theta$ | cut off emigration function | $3 \cdot \sigma_{a_S}$ |
| $b$ | shape parameter of the emigration function | (0,19) (cons.)<br>(-20,19) (plants) |
| | **Output variables** | |
| $\overline{\tau}$ | mean distance between all habitat patches, with $\tau_{nm}$, the absolute distance between patches $n$ and $m$, and $(Z^2 - Z)$, the total number of potential directed links between all $Z$ habitat patches | $\frac{\sum_{n,m=1}^{Z} \tau_{nm}}{Z^2-Z}$ |
| $\rho_i$ | landscape connectance of species i, with $L_i$, the number of directed dispersal links of species $i$ | $\frac{L_i}{Z^2-Z}$ |

### S4 Numerical simulations and data analysis

We constructed 30 model food webs, each comprising 10 plant and 30 animal species. To avoid confounding effects of different initial species diversities, we kept both the number of species $S$ and the fraction of plants and animals constant among all food webs. For each simulation, we randomly generated a landscape of size $Q$ (edge length of a square landscape) with $Z$ randomly distributed habitat patches. To test each food web across a gradient of number of habitat patches and habitat isolation, we drew the number of habitat patches, $Z$, from the inclusive interval $(10, 69)$ and the size of the landscape, $Q$, from the inclusive interval $(0.01, 10)$ using a stratified random sampling approach (see also section S2 for further information). With this approach, we generated landscapes on two independent gradients covering two aspects of fragmentation, namely number of fragments and habitat isolation. To cover the full parameter range of $Z$ and $Q$, we simulated each food web on 72 landscapes resulting in a total of 2160 simulations. We achieved a full range for the gradient of habitat isolation (landscape connectance ranging from 0 to 1, figure S3c). The upper limit for the number of patches was chosen to conform to the maximum usage time of 10 days per simulation on the high-performance-cluster we used [24]. Additionally, we performed dedicated simulation runs to reference the two extreme cases, i.e. (1) landscapes in which all patches are direct neighbours without a hostile matrix, and thus, no dispersal mortality, and (2) fully isolated landscapes, in which no species can bridge between patches, and thus, a dispersal mortality of 100% .

For each simulation run, we initialised our model with random conditions: Each habitat patch $z$ holds a random selection of 21 to 40 species (with each of the 40 species of the full food web existing on at least one patch) and initial biomass densities $B_{i,z}$ and nutrient concentrations $N_l$ ($l \in 1, 2$) were randomly sampled with uniform probability density within the intervals $(0, 10)$ for $B_{i,z}$ and $(S_l/2, S_l)$ for $N_l$, respectively. Here, $S_l$ are the supply concentrations of the nutrients, which are constant on all habitat patches but differ between the two nutrients. See table S2 and Schneider *et al.* [1] for further information on the nutrient dynamics.

Starting from these random initial conditions, we numerically simulated local food web and dispersal dynamics over 50,000 time steps by integrating the system of differential equations implemented in C++ using procedures of the SUNDIALS CVODE solver version 2.7.0 (backward differentiation formula with absolute and relative error tolerances of $10^{-10}$ [25]). Successful dispersal between local populations thereby enabled species to establish populations on patches where they were initially absent. For numerical reasons, a local population was considered extinct once $B_{i,z} < 10^{-20}$, and $B_{i,z}$ was then immediately set to 0.

#### Output variables

We recorded the following output variables for each simulation run: (1) the mean biomass density of each species $i$ on each habitat patch $z$ over the last 20,000 time steps to capture oscillations, $\overline{B}_{i,z}$; (2) the number of habitat patches in a landscape, $Z$; (3) habitat isolation, i.e. the mean distance between all habitat patches, $\overline{\tau}$ (see table S2); and (4) the landscape connectance of each species $i$, $\rho_i$ (see table S2). Thus, $\rho_i$ determines the ability of a species to connect habitat patches in a fragmented landscape.

**Statistical models and data visualisation** We tested for correlation between initialised and emerged $\beta$-diversity, which was however not the case (see section S9). Further, we used generalised additive mixed models (GAMM) from the mgcv package in R [26] to visualise the impact of number of patches and habitat isolation on species diversity. To fit the model assumptions, we logit-transformed $\overline{\alpha}$-diversity, and log-transformed $\beta$-diversity. We analysed each diversity index separately, with the number of patches $Z$ (log-transformed), the mean patch distance $\overline{\tau}$ (log-transformed) and their interaction as fixed effects and the ID of the food web (1 - 30) as random factor (with normal distribution for $\overline{\alpha}$- and $\beta$-diversity, and binomial distribution for $\gamma$-diversity). Similarly, we analysed the mean biomass densities, $\overline{B}_{i,z}$ (log-transformed), and species-specific landscape connectance, $\rho_i$, for each species (ID 1 - 40) using GAMM with a normal distribution. We used the mean patch distance, $\overline{\tau}$, as fixed effect and the food web ID (1 - 30) as random effect.

#### Analysis

Out of the 2160 simulations we started, 57 were terminated by reaching the maximum usage time of 10 days per simulation on the high-performance-cluster we used [24]. We further deleted 30 simulations as they had entirely isolated landscapes with no dispersal links. We performed all statistical analyses in R version 3.3.2. [27] using the output of the remaining 2073 simulations. See also section S8 for additional information.

**Species diversity** We quantified Whittaker's $\alpha$-, $\beta$-, and $\gamma$-diversity [28] using presence-absence data derived from the recorded mean biomass densities, $\overline{B}_{i,z}$, counting species $i$ present on patch $z$ when $\overline{B}_{i,z} > 10^{-20}$. In Whittaker's approach, $\alpha$ accounts for the local species richness, $\beta$ is the component of regional diversity that accumulates from compositional differences between local communities, and $\gamma$ is the regional diversity, i.e. the species richness at the landscape scale [28]. We relate $\alpha$, $\beta$ and $\gamma$ to each other using multiplicative partitioning [28], i.e. $\alpha \cdot \beta = \gamma$. Here, we use $\alpha$ averaged over all habitat patches $Z$ (which we hereafter refer to as $\overline{\alpha}$) to get a measure at the landscape level comparable to $\beta$ and $\gamma$.

#### S5 Maximum trophic level

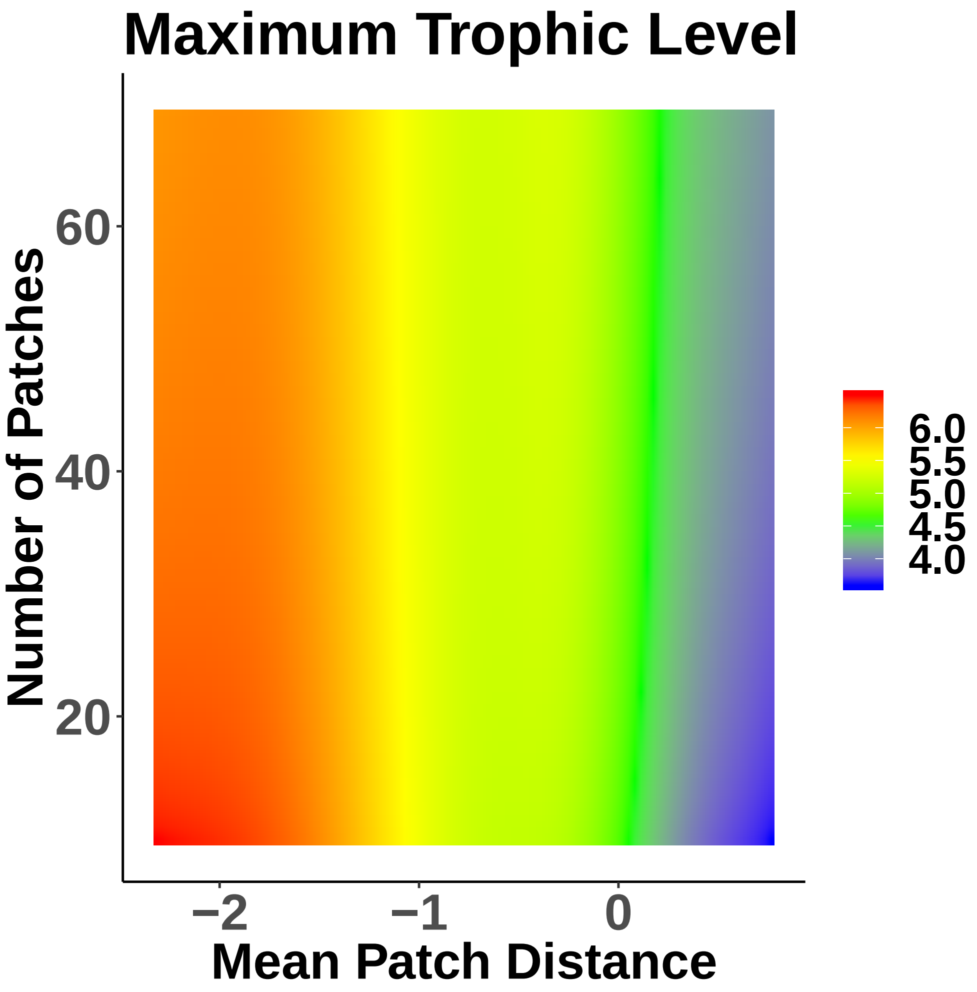


Figure S2: Heatmap visualising the maximum trophic level within a food web (colour-coded; z-axis) in response to habitat isolation, i.e. the mean patch distance ($\overline{\tau}$, $\log_{10}$-transformed; x-axis) and the number of habitat patches (Z; y-axis). The heatmap was generated based on the statistical model predictions (see the methods section in the manuscript). The loss of species diversity driven by habitat isolation also translates into a loss of the maximum trophic level.

### S6 Additional simulations with a constant maximum dispersal distance

We repeated all simulations with a constant maximum dispersal range for all species of $\delta_{const.} = 0.5$, i.e. all species have the same spatial network, to understand the effect of the dispersal advantage of larger animals. The results from these simulations are very similar to the results with the species-specific scaling of dispersal ranges, showing the same biomass density drop of larger animals at low mean distances (figure S3).

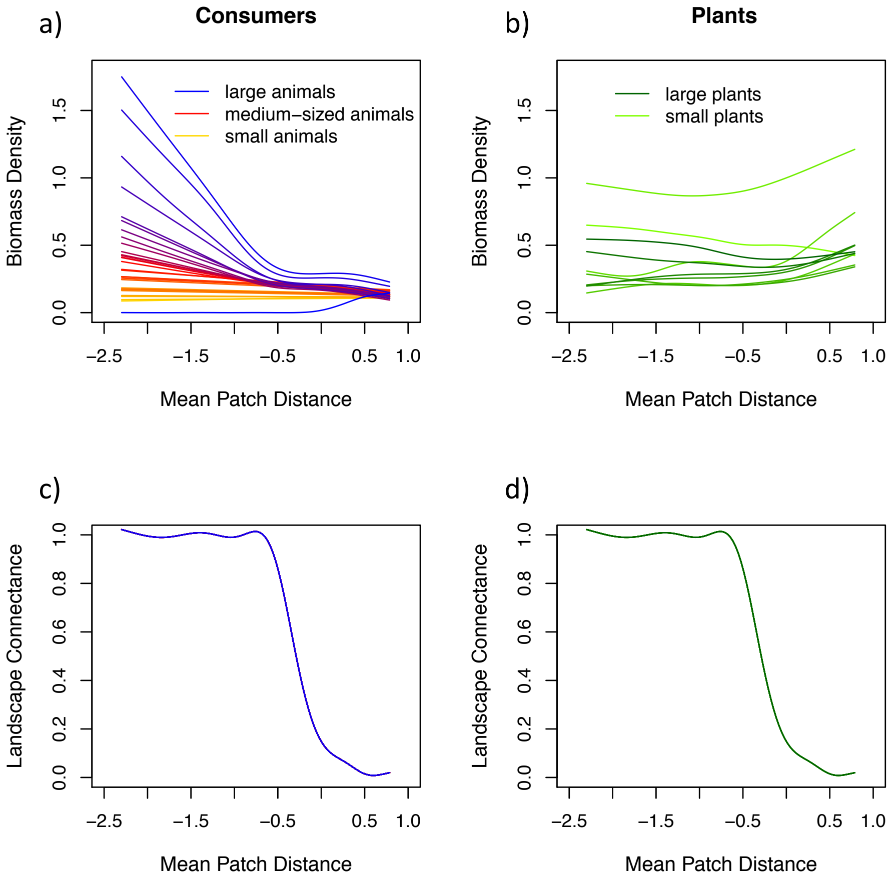


Figure S3: Top row: Mean biomass densities of consumer (a)and plant species (b) over all food webs ($B_i$, $\log_{10}$-transformed; y-axis) in response to habitat isolation, i.e. the mean patch distance ($\overline{\tau}$, $\log_{10}$-transformed; x-axis). Each colour depicts the biomass density of species $i$ averaged over all food webs: (a) colour gradient where orange represents the smallest, red the intermediate and blue the largest consumer species; (b) colour gradient where light green represents the smallest and dark green the largest plant species. Bottom row: Mean species-specific landscape connectance ($\rho_i$; y-axis) for consumer species (c) and plant species (d) over all food webs as a function of the mean patch distance ($\overline{\tau}$, $\log_{10}$-transformed; x-axis), using the same maximum dispersal distance for all species, $\delta_{const} = 0.5$.

### S7 Additional simulations of the two extreme cases

To explore the extreme cases of fragmentation in our model framework, we conducted additional simulations with emigration but no immigration on patches to represent completely isolated patches (disconnected), and landscapes with patches containing all species of the meta-food-web and neither emigration nor immigration to represent one joint landscape with no fragmentation (joint). For the disconnected scenario we simulated 12 replicates for each of the 30 food webs covering in the same stratified random gradient of patch numbers between 10 and 69 as in the main simulations and were also initialised with a subset of species (see the methods section in the paper). For the joint scenario we simulated 20 replicates for each food web containing 2 independent patches initialised with all species and no dispersal.

**(1) Joint scenario with no dispersal mortality** $\overline{\alpha}$-diversity is on average 37.621, $\gamma$-diversity 37.172 and $\beta$-diversity 1.004 (figure S4, purple triangle).

**(2) Fully isolated scenario with 100% dispersal mortality** $\overline{\alpha}$-diversity is on average 11.945, $\gamma$-diversity 32.801 and $\beta$-diversity 2.876 (figure S4, orange triangle).

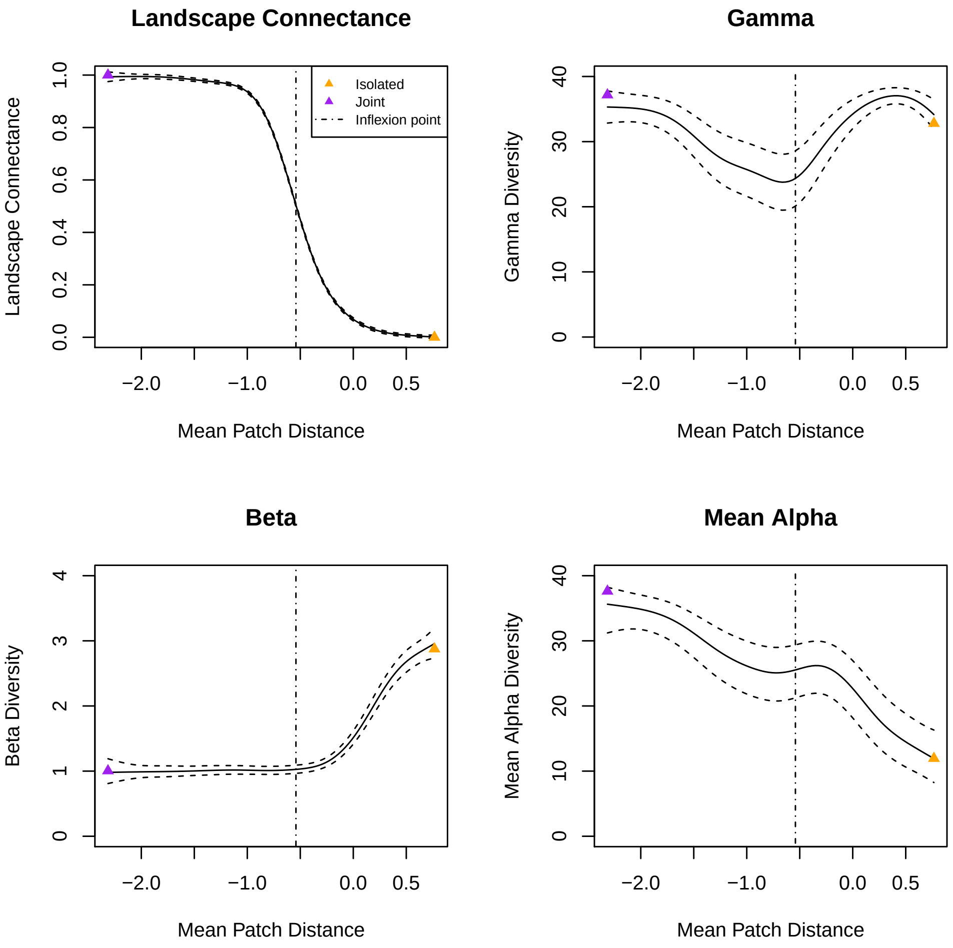


Figure S4: Shown are model predictions for landscapes with 40 patches across the whole gradient of the mean patch distance ($\overline{\tau}$, $\log_{10}$-transformed; x-axis). Top-left panel showing the landscape connectance averaged over all species (y-axis) as response to the mean patch distance ($\overline{\tau}$, $\log_{10}$-transformed; x-axis). Subsequent panels showing $\gamma$-diversity, $\beta$-diversity and $\overline{\alpha}$-diversity (y-axes) in response to the mean patch distance ($\overline{\tau}$, $\log_{10}$-transformed; x-axis). Purple triangles represent reference points from dedicated simulations in a joint scenario and orange triangles for fully isolated scenarios (see section S7).

### S8 Sensitivity analysis

We tested the effect of randomly drawn dispersal parameters (maximum dispersal rate, $a$, and the shape of the dispersal function, $b$; see the manuscript, equation 3) on mean $\alpha$-, $\beta$- and $\gamma$-diversity for consumers and plants respectively. We used generalised additive mixed models (GAMM) from the mgcv package in R for all sensitivity analyses. To fit the model assumptions, we logit-transformed $\overline{\alpha}$-diversity, and log-transformed $\beta$- and $\gamma$-diversity.The emigration parameters were separately used as fixed effects and the ID of the food web (1 - 30) as random factor (with normal distribution for $\overline{\alpha}$- and $\beta$-diversity, and binomial distribution for $\gamma$-diversity). Both parameters show no strong effect in all tested cases (figure S5 - S7). Only the maximum emigration rate $a$ of consumers shows a small negative effect on $\overline{\alpha}$-diversity (figure S5). As a higher maximum emigration rate results in an overall larger loss therm due to dispersal, which fits to our general findings.

Additional sensitivity analysis for interference competition, allometric exponent for attack rates of consumer species, exponents for handling time, hill coefficient and nutrient turnover rate were omitted as they were tested thoroughly in [1]. There, the dynamics of the food web model were shown to be robust to changes in model parameters. For each of the 2073 simulation runs the parameters of the trophic interactions were independently sampled from appropriate probability distributions within ecologically reasonable limits (see table 1). To account for the stochastic nature of the algorithm provided by Schneider *et al.* [1] by which food web topologies are created, we generated an ensemble of 30 food webs by randomly sampling 30 sets of species body masses.

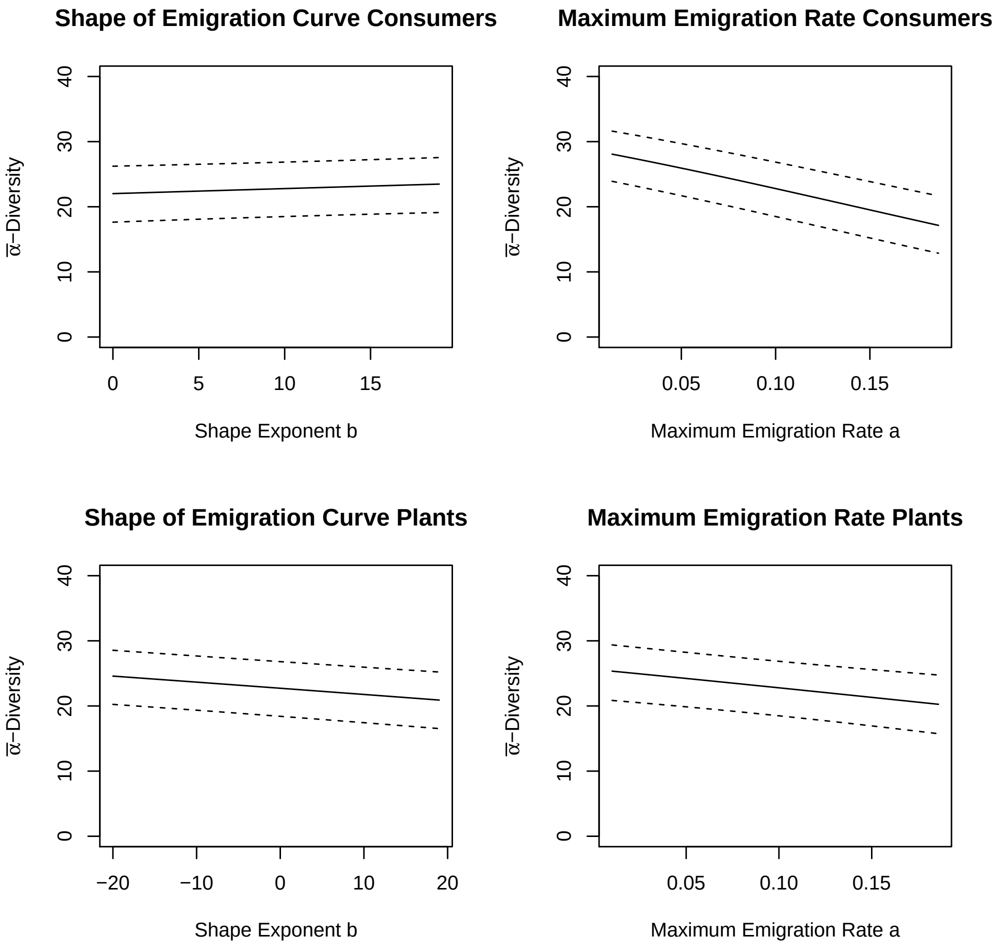


Figure S5: $\overline{\alpha}$-diversity (y-axes) of consumers and plants in dependence of the maximum emigration rate, $a$, and the shape of the emigration function, $b$ respectively (x-axes).

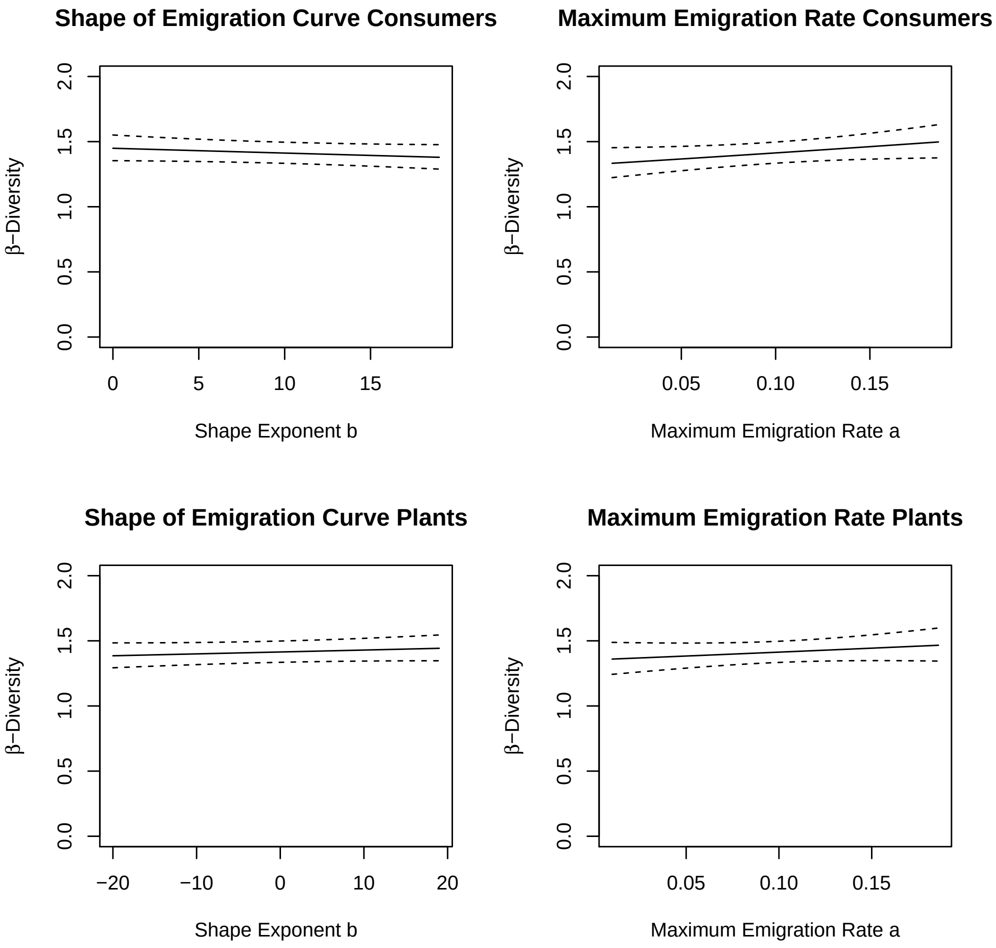


Figure S6: $\beta$-diversity (y-axes) of consumers and plants in dependence of the maximum emigration rate, $a$, and the shape of the emigration function, $b$ respectively (x-axes).

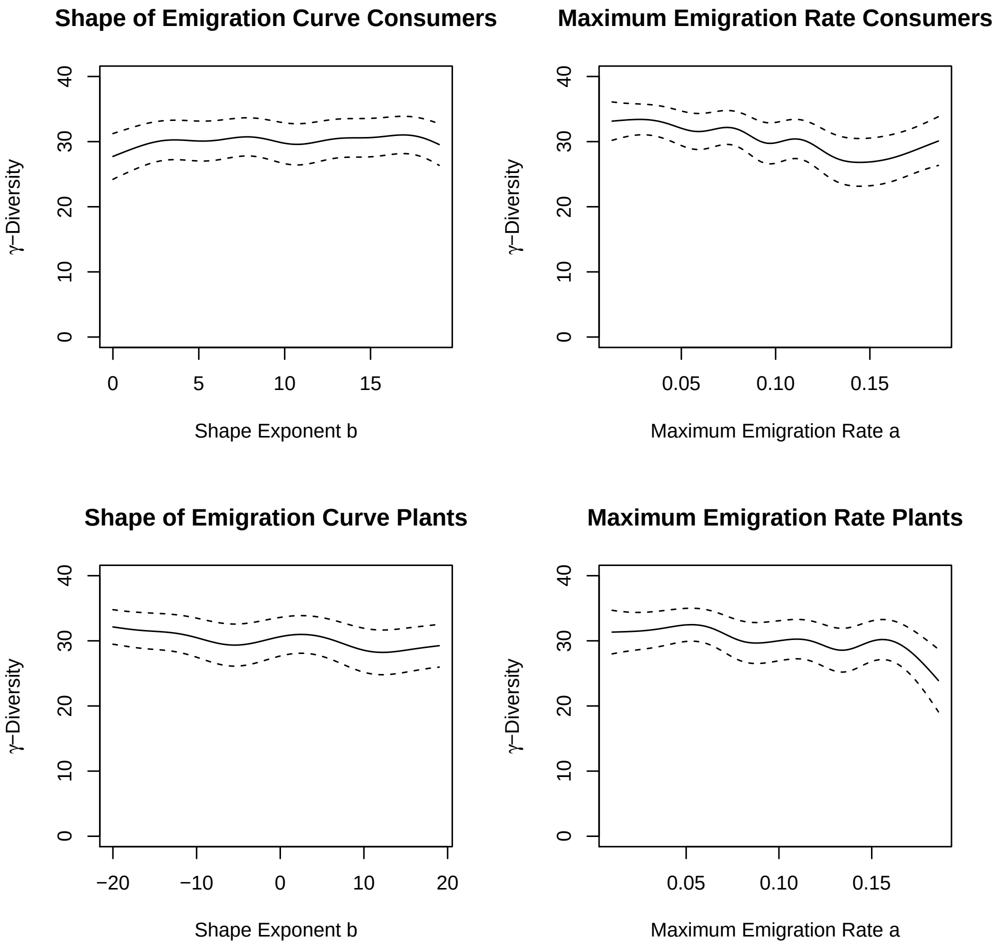


Figure S7: $\gamma-$diversity (y-axes) of consumers and plants in dependence of the maximum emigration rate, $a$, and the shape of the emigration function, $b$ respectively (x-axes).

### S9 Initial and post-simulation $\beta$-diversity

To see how the initialised $\beta$-diversity (see section S4) influenced the post-simulation $\beta$-diversity we performed a generalised additive mixed model (GAMM) from the mgcv package in R with the initial $\beta$-diversity as fixed effect and the post-simulation $\beta$-diversity as the response variable. Both were log-transformed to fit model assumptions. The post-simulation $\beta$-diversity and initial $\beta$-diversity were not correlated. This suggests that the initial $\beta$-diversity which is due to initialising the patches in the landscape with only a subset of species from the regional species pool does not

194 influence the post-simulation $\beta$-diversity delectably (approximate p-value: 0.518) (figure S8).

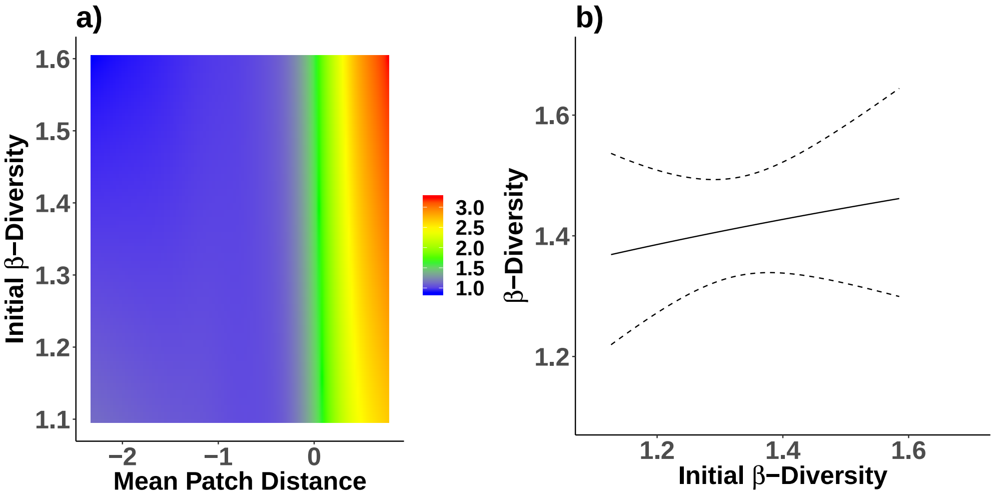


Figure S8: (a) The post-simulation $\beta$-diversity (y-axis) and the initial $\beta$-diversity (x-axis) were not correlated. (b) Heatmap visualising $\beta$-diversity (colour-coded; z-axis) in response to habitat isolation, i.e. the mean patch distance ($\overline{\tau}$, $\log_{10}$-transformed; x-axis) and the initial $\beta$-diversity (y-axis). The heatmap was generated based on the statistical model predictions (see the methods section in the manuscript). In strongly isolated landscapes $\beta$-diversity increases slightly with higher initial $\beta$-diversity. However, post-simulation $\beta$-diversity is higher than the initial $\beta$-diversity.

### S10 Standard errors in biomass densities

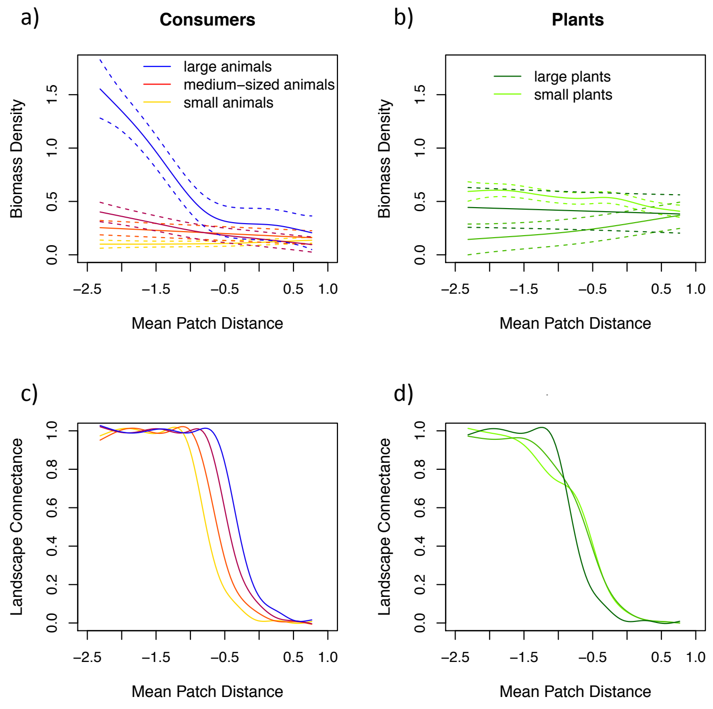


Figure S9: Top row: Mean biomass densities [log10(biomass density -1)] with standard errors [± 2*SE] for four exemplary animal consumer species (a) and three exemplary basal plant species (b) over all food webs ($B_i$, $\log_{10}$-transformed; y-axis) in response to habitat isolation, i.e. the mean patch distance ($\overline{\tau}$, $\log_{10}$-transformed; x-axis). Each colour depicts the biomass density of species $i$ averaged over all food webs: (a) colour gradient where orange represents the smallest, red the intermediate and blue the largest consumer species; (b) colour gradient where light green represents the smallest and dark green the largest plant species. Bottom row: Mean species-specific landscape connectance ($\rho_i$; y-axis) for consumer (c) and plant species (d) over all food webs as a function of the mean patch distance ($\overline{\tau}$, $\log_{10}$-transformed; x-axis).
